# A synthetic biology approach to assemble and reboot clinically-relevant *Pseudomonas aeruginosa* tailed phages

**DOI:** 10.1101/2023.06.23.546310

**Authors:** Thomas IPOUTCHA, Ratanachat RACHARAKS, Stefanie HUTTELMAIER, Cole WILSON, Egon A OZER, Erica M HARTMANN

## Abstract

The rise in frequency of antibiotic resistance has made bacterial infections, specifically *Pseudomonas aeruginosa*, a cause for greater concern. Phage therapy is a promising solution that uses naturally isolated phages to treat bacterial infections. Ecological limitations, which stipulate a discrete host range and the inevitable evolution of resistance, may be overcome through a better understanding of phage biology and the utilization of engineered phages. In this study, we developed a synthetic biology approach to construct tailed phages that naturally target clinically-relevant strains of *Pseudomonas aeruginosa*. As proof of concept, we successfully cloned and assembled the JG024 and DMS3 phage genomes in yeast using transformation-associated recombination (TAR) cloning and rebooted these two phage genomes in two different strains of *P. aeruginosa*. We identified factors that affected phage reboot efficiency like the phage species or the presence of antiviral defense systems in the bacterial strain. We have successfully extended this method to two other phage species and observed that the method enables the reboot of phages that are naturally unable to infect the strain used for reboot. This research represents a critical step towards the construction of clinically-relevant, engineered *P. aeruginosa* phages.

**Importance:** *Pseudomonas aeruginosa* is a bacterium responsible for severe infections and a common major complication in cystic fibrosis. The use of antibiotics to treat bacterial infections has become increasingly difficult as antibiotic resistance has become more prevalent. Phage therapy is an alternative solution that is already being used in some European countries, but its use is limited by narrow host range due to the phage receptor specificity, the presence of antiviral defense systems in the bacterial strain, and the possible emergence of phage resistance. In this study, we demonstrate the use of a synthetic biology approach to construct and reboot clinically-relevant *P. aeruginosa* tailed phages. This method enables a significant expansion of possibilities through the construction of engineered phages for therapy applications.

## Introduction

*Pseudomonas aeruginosa* (PA) is a Gram-negative bacterium responsible for 51,000 infections with 2,700 deaths in the US every year ^1^ and approximately 559,000 deaths globally in 2019 ^2^. PA is also a common complication of cystic fibrosis (CF), with 80% of CF patients developing PA infection ^3^ and causing chronic infection in 41% of un-transplanted adults with CF ^4^. Antimicrobial resistance of PA infections has become an increased concern ^1, 5^. This is particularly the case in low-and middle-income countries where multidrug-resistant bacteria are more prevalent ^6^.

Phage therapy is a promising alternative for treating infections ^7–10^. In 2022 the World Health Organization included it as a priority to fight antibiotic resistance, which is classified as a major concern over the next 5-10 years ^11^. For PA, phage therapy with naturally isolated phages has been developed successfully^7^; but the use of phage is limited by the phage specificity, which depends on the presence of phage receptor and defense systems (e.g., CRISPR systems, Restriction-Modification (RM)). Furthermore, even in sensitive strains, resistance appears through phage receptor mutations ^12^. To avoid resistance, alternatives like phage cocktails and/or combinations of phages and antibiotics have been used ^13, 14^. Unfortunately, not all combinations are synergistic ^15^ and a greater understanding of phage-bacteria interactions is needed to choose optimal combinations.

Phage engineering has the potential to improve phage therapy efficiency and avoid phage resistance^16^. The intent is to design phage therapy specific to the bacterial strain considering the phage receptor and the presence of antiviral defense systems to make the application of phage safer and more effective. Phage engineering encompasses a variety of applications, including inhibiting replication and changing the cargo carried. For example, phagemids, which consist of a phage capsid carrying a plasmid, cannot replicate in nature, have been designed in response to the need for safe technology. Engineered phagemids have been used to deliver a CRISPR system to antimicrobial-resistant strains of *Staphylococcus aureus* ^17^ or deliver antimicrobial enzymes ^18^. While this approach is promising, it is restricted to well-characterized phage like M13 in *Escherichia coli* or P1 in PA ^19^.

Phages can also be modified to be more suitable for therapeutic applications, e.g., to change phage host range by altering the phage tail fiber ^20^, or adding anti-CRISPR to bypass adaptative defense systems ^21^. These modifications are often performed using homologous recombination in the host bacteria ^10^. However, those methods are restricted to small rearrangements of non-essential proteins and are limited by the recombination efficiency ^22^. Other platforms have been used for both phage construction and reboot. For example, Cheng *et al.* ^23^ used *E. coli* to assemble, edit, and reboot a large panel of phages, including PA phages, to target Gram-negative bacteria, but as acknowledged in the study, no clinically relevant tailed phage have been rebooted and the methodology does not work for all phages. This limitation has been discussed in several papers ^24, 25^ and could be explained by the presence of toxic proteins encoded in the phage genome and subsequently expressed in *E. coli* ^26, 27^.

To avoid the limitations associated with working in *E. coli*, it is possible to separate phage engineering into two steps: 1) assembly of the synthetic genome and 2) production of phage particles with a synthetic genome (reboot). One well-known platform for construction and engineering of various bacterial and viral genomes is the yeast *Saccharomyces cerevisiae* ^28–30^. In contrast to *E. coli*, prokaryotic DNA, including toxic molecules that could be encoded by phages, is rarely expressed in yeast and does not impact yeast fitness ^31^. Yeast has been used for this purpose to clone or construct synthetic phage genomes, changing tail fiber specificity ^25^. While yeast is useful for producing synthetic phage genomes, they are incapable of producing phage particles, i.e., performing “reboot.” For Gram-negative phages, reboot is still performed in *E. coli*, which again restricts the method to only certain phages. Recently, some *S. aureus* and *Enterococcus faecalis* phages were constructed in yeast and rebooted directly in *S. aureus* ^24^. Furthermore, *Pseudomonas* phage vB_PaeP_PE3 has been cloned and engineered in yeast to construct a reduced phage genome, which was successfully rebooted in PAO1^32^. Although vB_PaeP_PE3 is part of the *Autographiviridae* family and cannot infect clinically relevant PA strains ^32, 33^, this study demonstrates the feasibility of genome manipulation in yeast. It remains, however, unclear how generalizable the results are and whether all PA phage are amenable to this process.

Engineering phage genomes in yeast enables large and diverse modifications, but the resulting genomes still need to be rebooted. In the current study, we examine the use of yeast for genome engineering, followed by reboot using PA. We focus on addressing limitations in the reboot process by examining JG024, a member of the genus *Pbunavirus*, which are lytic phage that infect numerous clinically relevant PA strains and are thus considered candidates for phage therapy ^34–38^. JG024 ^39^ was extensively studied for this application in combination with antibiotics ^40^. We develop a methodology for construction of synthetic phage particles using transformation-associated recombination (TAR)-cloning with yeast followed by rebooting the phage DNA into *P. aeruginosa* to produce viable phage particles (Figure 1). Comparing reboot success between different phages in PA led us to identify factors that limit phage reboot, like phage DNA circularity, phage-specific characteristics, and host antiviral defense systems. This work represents the first time PA phage of high interest for phage therapy applications are successfully rebooted from synthetic genomes produced in yeast.

**Figure 1:**
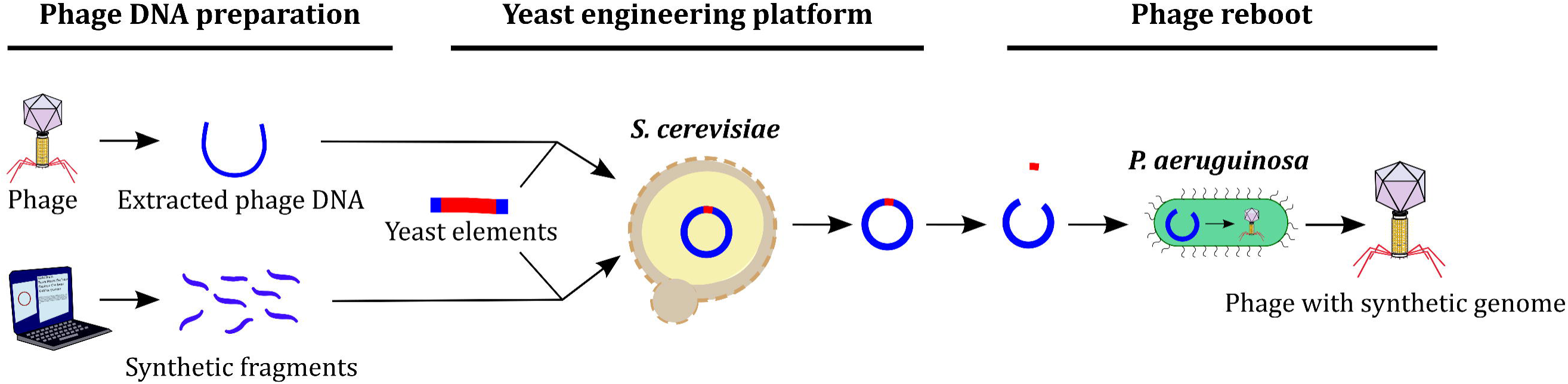
Schematic representation of the experimental procedure. Using direct extraction of the phage genome or the construction of overlapping fragments amplified by PCR, we were able to clone or construct the phage genome in yeast and maintain it using yeast elements. Next, extraction of yeast DNA and digestion by restriction enzymes allowed us to obtain full-length phage DNA that is free from yeast elements. Finally, PA transformation permitted us to obtain rebooted phage particles.

## Results

### Corroboration of a circular permuted JG024 genome

To enable the development of a successful cloning and reboot strategy, it is critical to characterize the genome of the phage in question. We thus sequenced the genome of our JG024 (Figure 2A), revealing both conserved structural features and population-level heterogeneity. Compared to the published JG024 genome (66,275 bp) ^39^, we observed two insertions, one G at position 29,132 (in 52% of short reads) and one A at position 55,007 (in 97% of short reads, 337^th^ amino acid position of ORF F358_gp71). We confirmed these two mutations by Sanger sequencing, indicating that they are not artefacts of the sequencing process but rather reflect population-level heterogeneity in the phage. Hybrid assembly using Unicycler, Trycycler or Flye generated a circular assembly of 66,277 bp (Figure 2A). In addition, Trycycler produced a linear assembly of 66,307 bp which was identified by CheckV to contain direct terminal repeats (DTRs) of 30 bp. PhageTerm was also assessed for the ability to use short-read sequencing data to identify the phage genome termini of the circular assembly ^41^. PhageTerm identified DTRs of 270 bp resulting in a linear genome of 66,547 bp. To verify the presence of either the 30 bp or 270 bp DTRs in the two linear assemblies, “primer walking” via Sanger sequencing was used with primers that anneal to the unique genomic site internal to the terminal repeats and directed to sequence across the repeats to the ends (Figure S1-A). For both linear assemblies, DTRs were not identified, as there was no termination of the sequence or decrease in signal intensity after the proposed DTR sequence. Instead, the sequence continued beyond the DTR suggesting a continuous sequence akin to a circular assembly. Only one known phage genome structure could result in circular assembly of phage dsDNA: circular permuted genomes. In this case, a packaging site (*pac* site) is usually recognized by a phage protein to initiate DNA packaging, but the terminase has poor specificity and nonspecific headful cleavage happens when the capsid is full, resulting in the presence of phage ranging from 98 to 110 % of the phage DNA molecule ^42^. Our assembly suggests that the JG024 genome is a circularly permuted genome and that the phage uses a headful packaging strategy.

**Figure 2:**
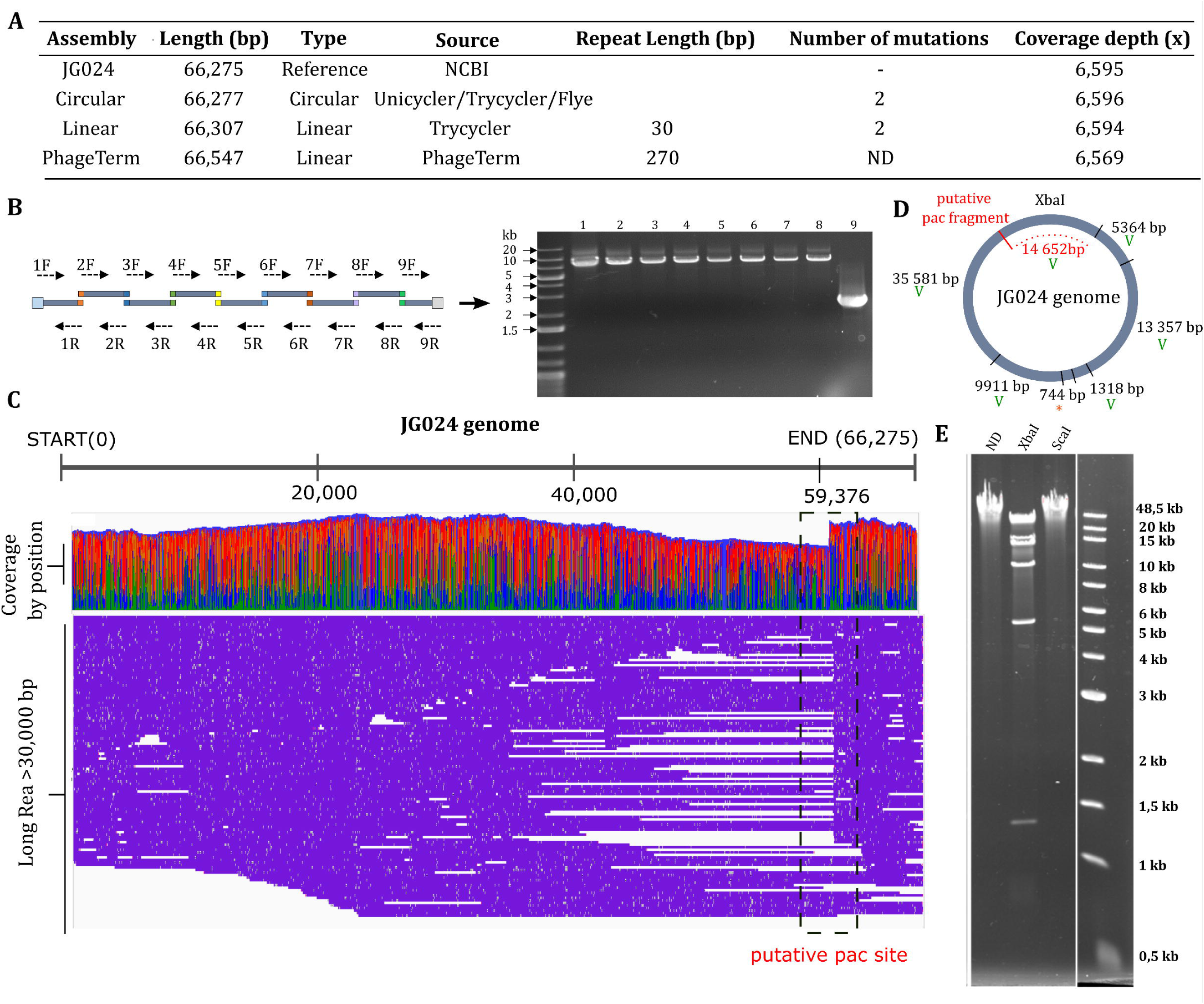
Analysis of JG024 genome. **A-** Overview of assembly results obtained using different software: Unicycler, Trycycler, Flye, and PhageTerm. **B-** DNA amplification of full JG024 genome with overlapping fragments. **C-** Visualization of mapped long reads >30,000 bp to JG024 reference genome by IGV software. Coverage by nucleotide position is also represented. **D-E-** Phage DNA digestion and migration on agarose gel and representation of attempted results. *ND*: undigested JG024 genome; *XbaI*: digestion of JG024 genome with XbaI; *ScaI*: digestion of JG024 genome with ScaI.

To corroborate this hypothesis experimentally, we successfully amplified the entire viral genome using primers to generate 9 overlapping fragments (Figure 2B). Although JG024 was previously identified to have a linear genome through exonuclease *Bal31* digestion ^39^, the amplification of the entire viral genome using overlapping fragments suggests that JG024 has no physical ends within the proposed assembly (Figure 2A). Furthermore, as each successfully amplified fragment must originate from at least some virion DNA molecules that contain the entire length of the fragment, this is consistent with the idea of a circularly permuted genome. Additionally, if we observe the global distribution of all long reads greater than 30,000 bp obtained from our sequencing efforts (Figure 2C), we observed a decrease of coverage depth between positions 50,000 and 60,000. We also see that a larger proportion (36/161) of reads start at position 59,376 (+/-5 bp). This could be the packaging series initiation site (*pac* sequence) recognized by the phage terminase protein for DNA packaging. As previously described for P22, SPP1 and P1 phages ^43^, restriction of circular permuted genomes results in fragments that would be predicted from a circular molecule, with an additional *pac* fragment sometimes observed. Concerning JG024, we observed that the genome was not sensitive to three enzymes (ScaI, NdeI, BsaI), suggesting that the DNA is methylated (Figure 2E). Using XbaI, we observed that the restriction digest profile corresponds to a circular permuted genome (Figure 2E, S1-B); this is in contrast to previous conclusions in the study by Garbe *et al*., which predicted that the genome was linear despite incongruous results from SacII digestion ^39^. Their conclusion was based on a linear map of the JG024 genome, but their result could correspond to a circular digestion profile (8.5 + 21.7 + 35.9 kb). In addition to the bands predicted from a circular assembly, we observed a restriction band around 15,000 bp that does not correspond to a band predicted from a linear profile. This band matches the predicted *pac* fragment starting from the putative *pac* site at position 59,376 bp (Figure 2C-2D-2E). Together, **these data suggest that the JG024 genome is circular permuted. This knowledge is important for designing the cloning strategy in yeast and will guide us to use linear-linear recombination to assemble and maintain the JG024 phage genome.**

#### Assessment of chloroform sensitivity and other parameters to improve reboot efficiency

To optimize the reboot protocol and avoid issues linked to low reboot or transformation efficiency, we assessed how different parameters affected phage titer. Chloroform is often used during phage production to destroy bacterial cells and release phage particles in the bacterial lysate ^44^. As chloroform affects 30% of tailed phages ^45^, we assessed the effect of chloroform on JG024. JG024 phage lysate was treated with chloroform before infecting PA14, and JG024 plaques were then enumerated using the double agar method. Chloroform significantly affected phage titer (*p*=0.004) which decreased 4.57-fold (78.2% reduction) compared to the untreated phage lysate (3.4 x 10^8^ PFU/mL) (Figure 3A), indicating that JG024 is sensitive to chloroform.

**Figure 3:**
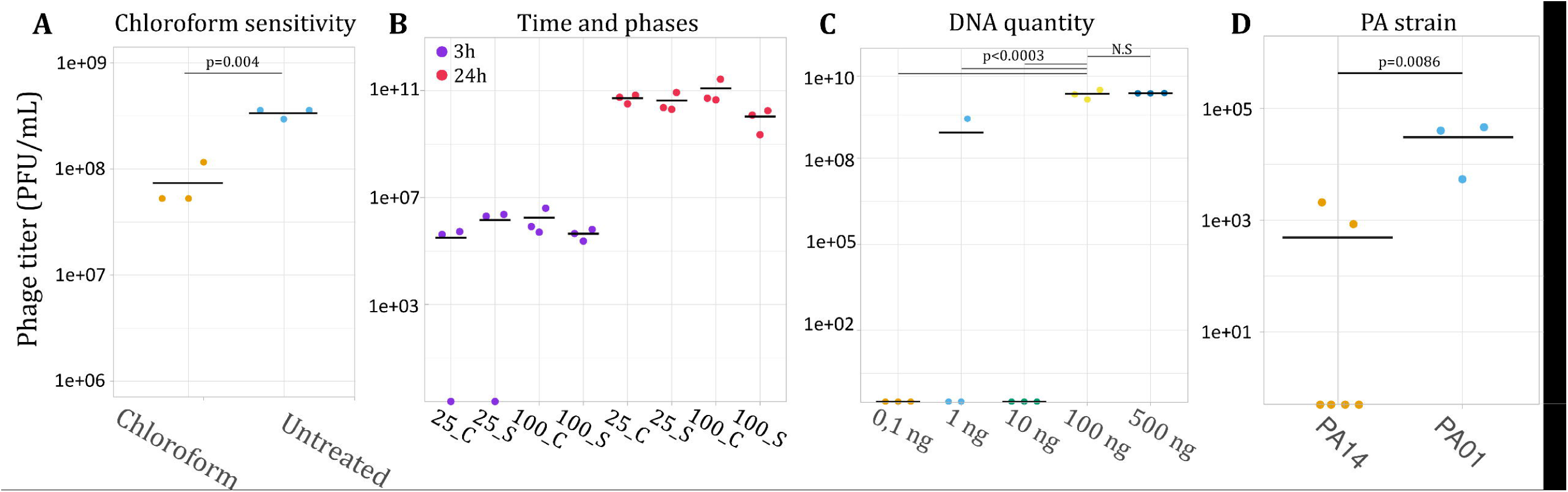
Optimization of the reboot protocol. Graphical representation for all parameters tested: **A**JG024 chloroform sensitivity, **B-** Recovery time (3 h or 24 h), phage phase (cell pellet: C, or supernatant: S) using 25 or 100 ng, **C-** DNA quantity using 0.1, 1, 10, 100 or 500 ng JG024 DNA, **D-** Effect of PA strain.

To investigate if JG024 phages are well released from PA14 cells during the rebooting process, JG024 gDNA (25 ng and 100 ng) was electroporated into electrocompetent PA14 cells and incubated for either 3 h or 24 h. After incubation, the cell suspension was pelleted and the supernatant was assessed directly for PFU to quantify the phages released naturally from phage-mediated cell lysis. The remaining cell pellet was washed three times with LB media, treated with chloroform, and assessed for PFUs to quantify the phages released primarily from chloroform treatment. The 3 h incubation was sufficient to observe PFUs and the longer recovery time significantly increased the number of PFUs (*p*=0.0001). This observation agrees with expectations for lytic phage in a sensitive bacterial culture. Furthermore, phage particles were found in the same quantity in the supernatant or bacterial pellet after chloroform release (Figure 3B).

We next attempted to reboot JG024 using different quantities of JG024 gDNA. The quantity of JG024 gDNA significantly affected phage titer (*p*=0.0003) as only gDNA quantities of at least 100 ng resulted in consistent PFUs. There was no significant difference in phage titer between 100 ng and 500 ng of gDNA (p>0.05) with phage titer reaching an average of 1.8 x 10^10^ PFU/mL for both DNA quantities (Figure 3C).

We finally investigated the effect of different PA strains on JG024 reboot efficiency. Using strain PAO1, we obtained a greater number of PFUs and more consistent results compared to PA14 (Figure 3D). These results indicate that specific host-strain factors are critical to phage infection and replication. Other transformation parameters, such as wash buffer (300 mM sucrose vs 1 mM MgSO4), MgSO4 concentration after electroporation (0 mM, 1 mM MgSO4, 10 mM MgSO4) and electroporation voltage (1.8 kV, 2.2 kV, 2.5 kV) were also tested (Figure S2). The buffer had a significant effect on the phage titer with the use of MgSO4 resulting in higher phage titer than sucrose (*p*=0.002). The phage titer from 2.2 kv was higher than the phage titer from 1.8 kv (*p*=0.02). **These data suggest that a reboot protocol without the use of chloroform can improve reboot efficiency. We also observed that a high concentration of JG024 phage DNA and PA strain-specific characteristics can increase reboot success.**

### Successful cloning and construction of JG024 genome in yeast

Based on previous work for the cloning of full bacterial and viral genomes ^28, 46–48^, we chose the yeast *S. cerevisiae* VL6-48N as a platform to clone and replicate JG024 DNA (Figure 4A). TAR-cloning ^49^ has been used extensively for the isolation and production of large genomic fragments from a variety of organisms.

**Figure 4:**
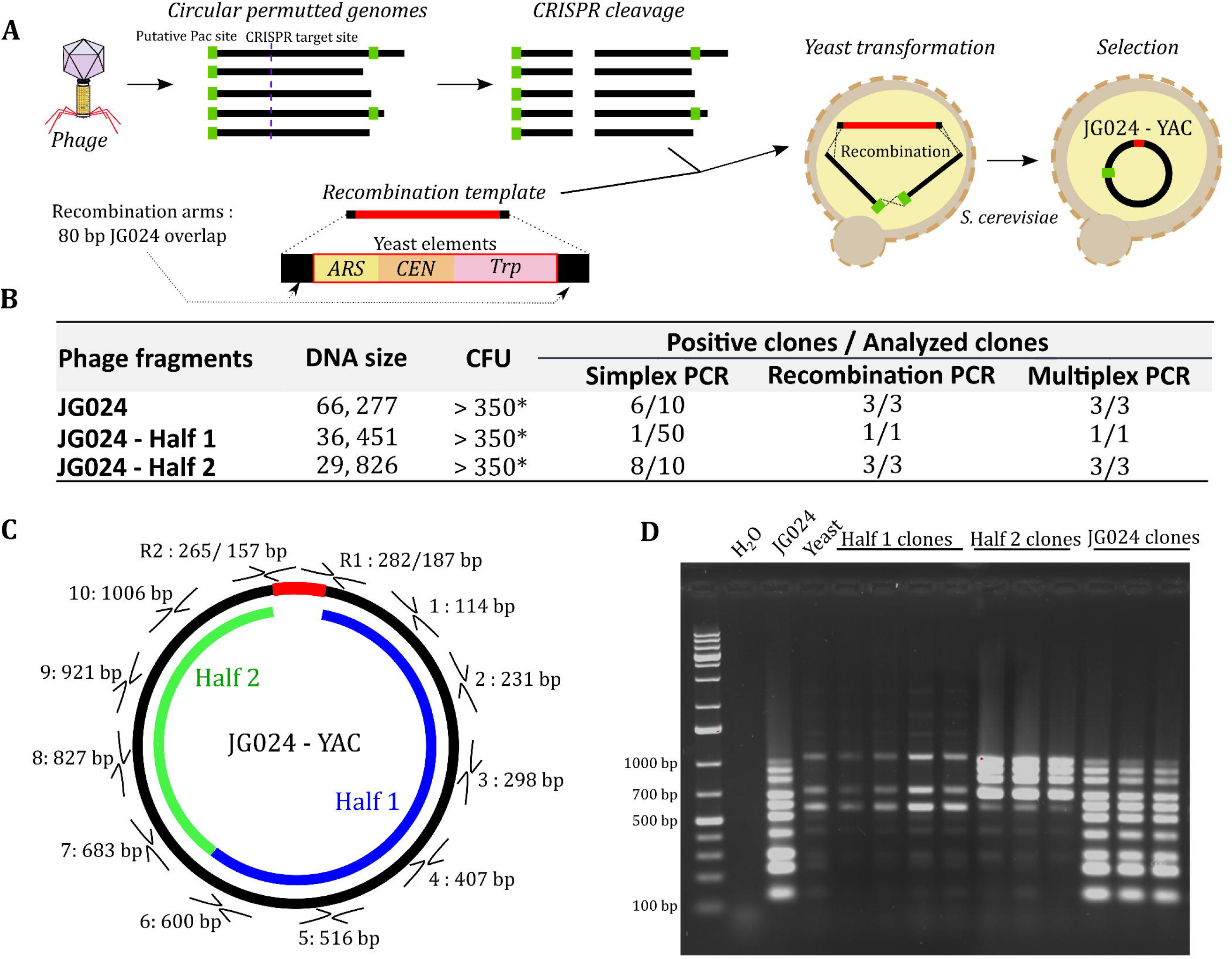
JG024 genome in yeast results by TAR-cloning. **A-** Schematic procedure of JG024 TAR-cloning. **B-** Recapitulative table of phage DNA cloning in yeast. Simplex PCR consists of one PCR that amplifies a single region of the genome. Recombination PCR involves amplification of recombination scars, and multiplex PCR uses a set of several primers to amplify multiple regions around the phage genome (in this case, 10). **C**Representation of the three batches of PCR done to validate phage genome in yeast. **D-** Example of multiplex PCR performed to validate phage genome integrity in clones.

For cloning JG024 in yeast, we used the full length JG024 genome and a recombination template flanked by 60 bp of homology (recombination arms), containing a centromeric sequence (CEN), autonomously replicating sequence (ARS), and an auxotrophic element for selection and maintenance in yeast (Trp) (Figure 4A). As we previously hypothesized that JG024 is circular permuted, terminal ends should be different on each copy of JG024 genome. We used in vitro cleavage using SpCas9-sgRNA, to target and cleave a precise location (target used: ACAATCCTCATAAGAAGTCGCGG) and obtain phage molecules linearized at the same position. After transformation, we obtained several hundred yeast colonies (Figure 4B) and screened 10 clones. We first validated the presence of phage DNA using a unique PCR amplifying 827 bp of the JG024 genome, and 6 yeast clones of the 10 screened showed amplification (Figure S3-A). We next validated the recombination event by amplifying recombination scars (Figure S3-B). Finally, for the presence of a full phage molecule, we performed multiplex PCR on ten JG024 parts (Figure 4D). Of 3 screened clones, all were validated as containing a circular JG024 genome. In addition to full size JG024 DNA, we also used simultaneously a second sgRNA to cut the genome in a second genome location (target used 2: CTAGTGTACGCTAGAATCAGTGG), and clone JG024 genome in two parts. We again used two recombination templates flanked by 60 bp of homology (recombination arms) specific for each JG024 fragments. For the first half, only 1 yeast clone out of 50 screened contained the expected phage DNA (Figure 4). In contrast, despite using the same JG024 DNA preparation for cloning and only different recombination arms, we obtained 8 clones of 10 screened that contained the second half. These results suggest that the TAR cloning efficiency is not equal for all genomes and that the recombination arms can impact the number of yeast clones to be screened.

To determine whether yeast strategy permits high throughput genomic manipulation, we attempted to synthetically reconstruct the JG024 genome from multiple PCR fragments. From phage DNA we amplified the JG024 genome in 3 overlapping DNA fragments (Figure 5A-5B). After transformation in yeast, we obtained 10 yeast colonies (Figure 5C), which is relatively low number of colonies compared to TAR-cloning (Figure 4B). However, as the assembly requires more recombination events than TAR-cloning, increasing the recombination arm’s length could improve the number of transformants. Despite this low colony number, we obtained 5/10 clones with full sized JG024 genomes. DNA stability over time is critical for maintaining and performing genome engineering in yeast. To test the stability of the synthetic JG024 genome in yeast, we performed 10 successive passages and observed the DNA integrity using multiplex PCR (Figure 4D). After 10 passages, we did not observe any DNA rearrangement and we thus concluded that JG024 phage DNA is stable in yeast. **In summary, we successfully cloned the JG024 genome in yeast directly from extracted phage genomic DNA. We further demonstrate the simultaneous use of two sgRNA for JG024 modification purposes. We also showed that synthetic DNA could be used for the construction of JG024 genomes with large DNA modifications.**

**Figure 5:**
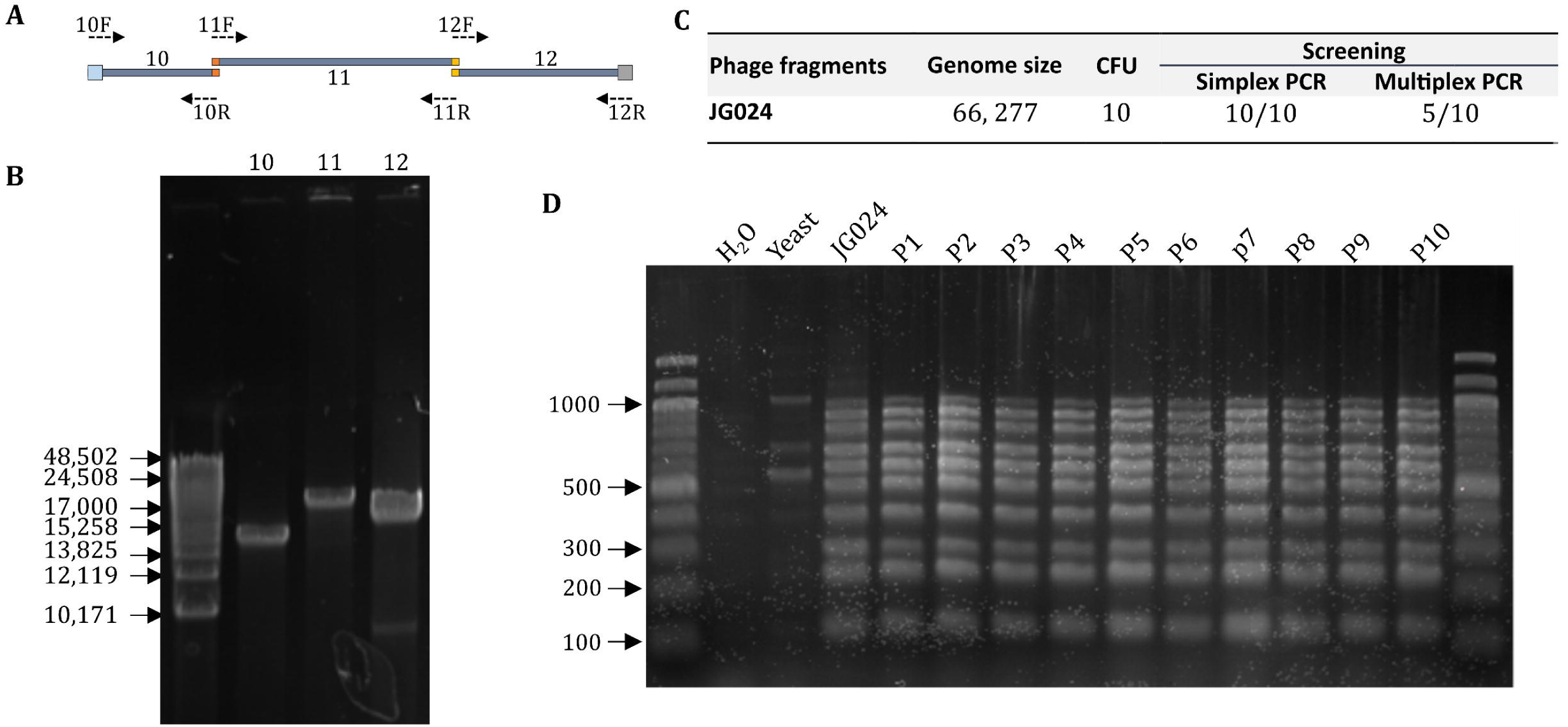
Construction of synthetic phage DNA in yeast. **A-** Three-fragment PCR design for JG024 genome amplification. **B-** PCR fragments obtained for JG024 cloning in yeast. **C-** Results obtained for assembly of JG024 fragments in yeast. **D-** JG024 stability results obtained by multiplex PCR on yeast clones during 10 passages.

### Unsuccessful cloning of JG024 and smaller fragments in *E. coli*

To enable downstream cloning in *E. coli*, we used a recombination template that contained not only the previously described yeast element but also an *E. coli* element (OriV, Chloramphenicol acetyl transferase gene). However, in three replicates, only 7, 0 and 0 *E. coli* colonies were obtained. Of those, only two were able to grow in liquid culture, and none showed the presence of JG024 DNA. We further attempted to clone the halved JG024 genome in *E.* coli, but no colonies were obtained for either half after 3 attempts at transformation. These results suggest that the size of the JG024 genome alone is not solely responsible for its toxicity in *E. coli*.

### Identification of phage-and host-specific limits to phage reboot

To reboot JG024 DNA from the yeast clones, we extracted DNA and first attempted to transform PA using 10 µg of yeast DNA extraction. However, no plaques were observed in either PA14 or PAO1 strains. We hypothesized that factors related to JG024 itself, bacterial factors in the strain that is used for rebooting or some combination of the two were inhibiting reboot of the synthetic JG024 construct.

To understand if the synthetic JG024 genomic construct itself is problematic for rebooting purposes, we attempted to replicate our observations with another phage. For comparison, we selected DMS3 ^50–52^, which is part of the Casadabanvirus family of phage. Similar to JG024 the genome is predicted to be circular permuted DNA ^53^. In addition, the DMS3 genome naturally encodes anti-CRISPR and anti-quorum sensing proteins ^53^. The genome of DMS3, at 36 kb, is also substantially smaller than that of JG024. We cloned DMS3 DNA in yeast using TAR-cloning and validated genome integrity as described for JG024 (Figure 6A). We next tried to transform synthetic DMS3 genomes in PA14 and PAO1. In contrast to JG024 (Figure 6B), we observed DMS3 plaques, but only in PAO1 strain. **We corroborate our previous findings that strain-level differences in hosts (PA14 or PAO1) impede or enhance reboot. Finally, we validate that DMS3 phage can be rebooted from a genome generated in yeast, which further suggests that phage-specific characteristics also impact the reboot success.**

**Figure 6:**
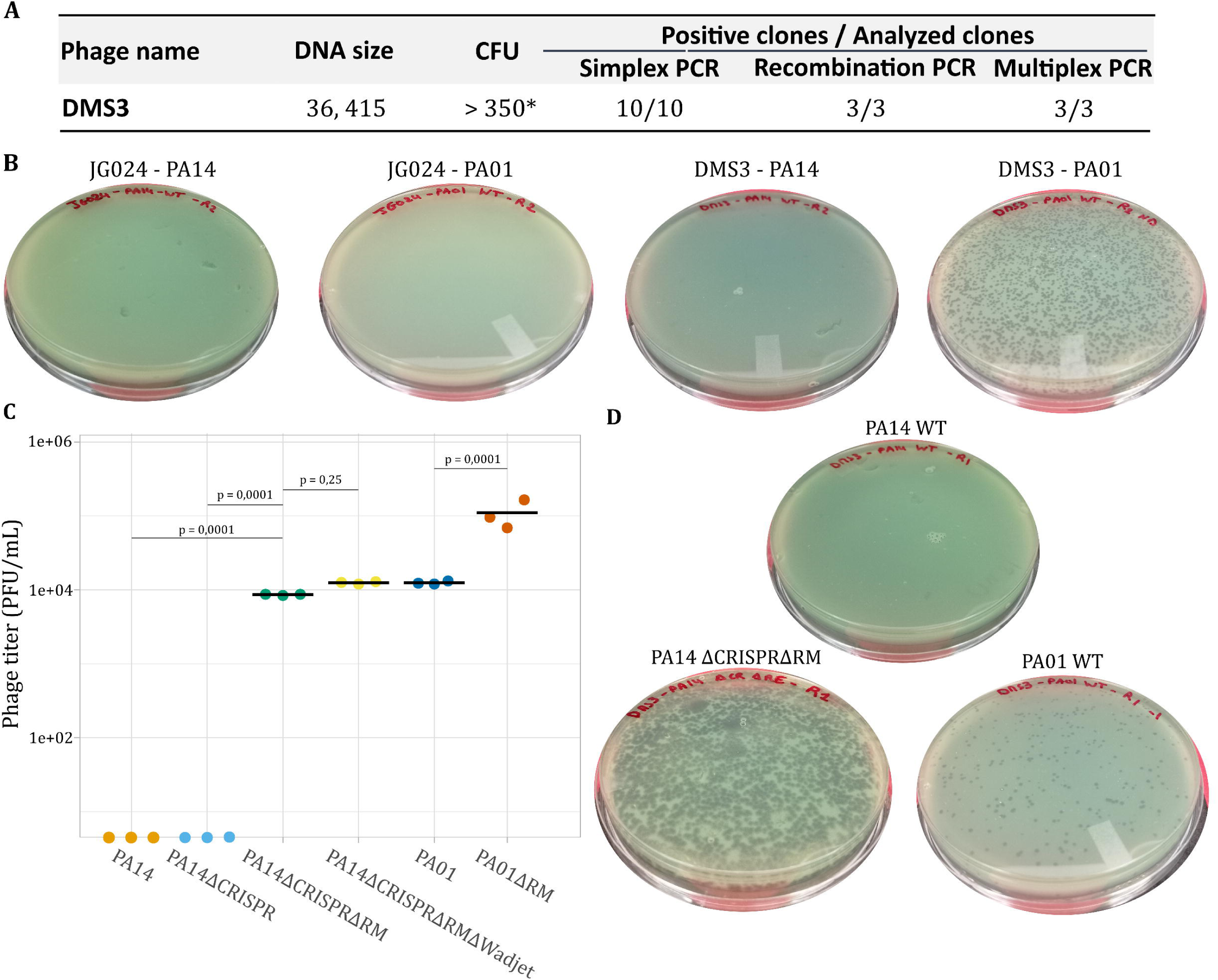
Comparison JG024 reboot and DMS3 and JG024 reboot in PA14 and PAO1. **A-** Table of in yeast cloning result for DMS3 genome. Simplex PCR consists of one PCR that amplifies a single region of the genome. Recombination PCR involves amplification of recombination scars, and multiplex PCR uses a set of several primers to amplify multiple regions around the phage genome (in this case, 6). **B-** Example reboot result from yeast DNA obtained for linearized DMS3 and JG024 genomes in PA14 and PAO1. **C-** Reboot of linear DMS3 phage DNA from yeast DNA in PA mutants. **D-** Example reboot result from yeast DNA obtained for linearized DMS3 genomes in PA14 and PAO1 mutants.

We next tried to understand why DMS3 phage can be rebooted in PA01, but not in PA14, and JG024 could not be rebooted in either. Multiple determinants are responsible for bacterial strain specificity, including differences in receptors, superinfection immunity or exclusion, and differences in antiviral defense systems ^54, 55^. As wildtype JG024 infects both PA14 and PAO1, we do not expect differences in expression of lipopolysaccharide (JG024 receptor) or superinfection to be major barriers to reboot. We thus hypothesized that the difference observed in reboot between PA14 and PAO1 could be linked to their antiviral defense systems. We identified these systems using PADLOC (Table S4) ^56^ and found at least 4 that could interact with DNA and impact reboot: a type-I restriction modification (RM) system in PAO1, and type-II RM, type I CRISPR and Wadjet systems in PA14. RM systems protect endogenous DNA and cleave exogenous DNA via methylation discrimination. Production of phage genome in yeast will affect its DNA methylation profile and could be a limitation to DNA transformation and phage reboot. Type I-F CRISPR system and Wadjet systems are composed of several proteins that possess nuclease activity and could then interact with phage exogenous DNA ^51, 57^, preventing DNA transformation and phage reboot. To determine if host antiviral systems were inhibiting phage rebooting from phage genome cloned in yeast, we used four PA mutant strains: PAO1ΔRE, PA14ΔCRISPR, PA14ΔCRISPRΔRE and PA14ΔCRISPRΔREΔWadjet, validated by whole genome sequencing (Table S3). Plasmid DNA transformation efficiency was similar between our WT strains and mutants (*p*>0.05) (Figure S5). Finally, we tried to reboot DMS3 phages using linear DNA from yeast extractions. As observed previously (Figure 6B), DMS3 reboot was not observed in either PA14 or PA14ΔCRISPR (Figure 6D). In contrast, when the type-II RM system was removed, we observed consistent reboot (*p*=0.0001) (Figure 6D). Additional removal of the Wadjet system resulted in a 1.4-fold increase in plaque numbers, however this improvement was not statistically significant (*p*=0.25). In PA01, reboot of the linearized synthetic DMS3 construct was previously successful in WT PAO1 (Figure 6B) but removal of the Type-I RM system resulted in an 8.8-fold increase in plaques (*p*=0.0001) (Figure 6D).

Next, we tried to reboot JG024 in the PA defense system knockouts. We did not obtain rebooted phage using PA14 or PA14ΔCRISPR (Figure 7) as observed for DMS3 phage, nor were we able to reboot using PAO1 as previously observed (Figure 6B). However, removing the Type-II RM system from PA14 enabled phage reboot, albeit only in 1 out of 3 replicates (Figure 7). When we removed the Wadjet system, we observed more consistent reboot with replicable results in comparison to the Type-II RM mutant (Figure 7). Finally, in PAO1, we also obtained reboot with JG024 DNA when the type-I RM was removed (Figure 7). If we compare JG024 and DMS3 reboot (Figure 6C-Figure 7), JG024 is less efficiently rebootable compared to DMS3 as we used 2.5 h of rebooting time for DMS3 instead of 6 h for JG024, but we still observed more efficient reboot for DMS3. **In summary, in addition to genome circularity (or removal of the yeast element), we observed that defense systems can interact with phage DNA produced in yeast. In particular, the PA14 type-II RM system can lead to total inhibition of phage reboot, whereas other systems may dampen efficiency or decrease repeatability. Finally, phage-defense system interactions are specific to the phage and host in question.**

**Figure 7:**
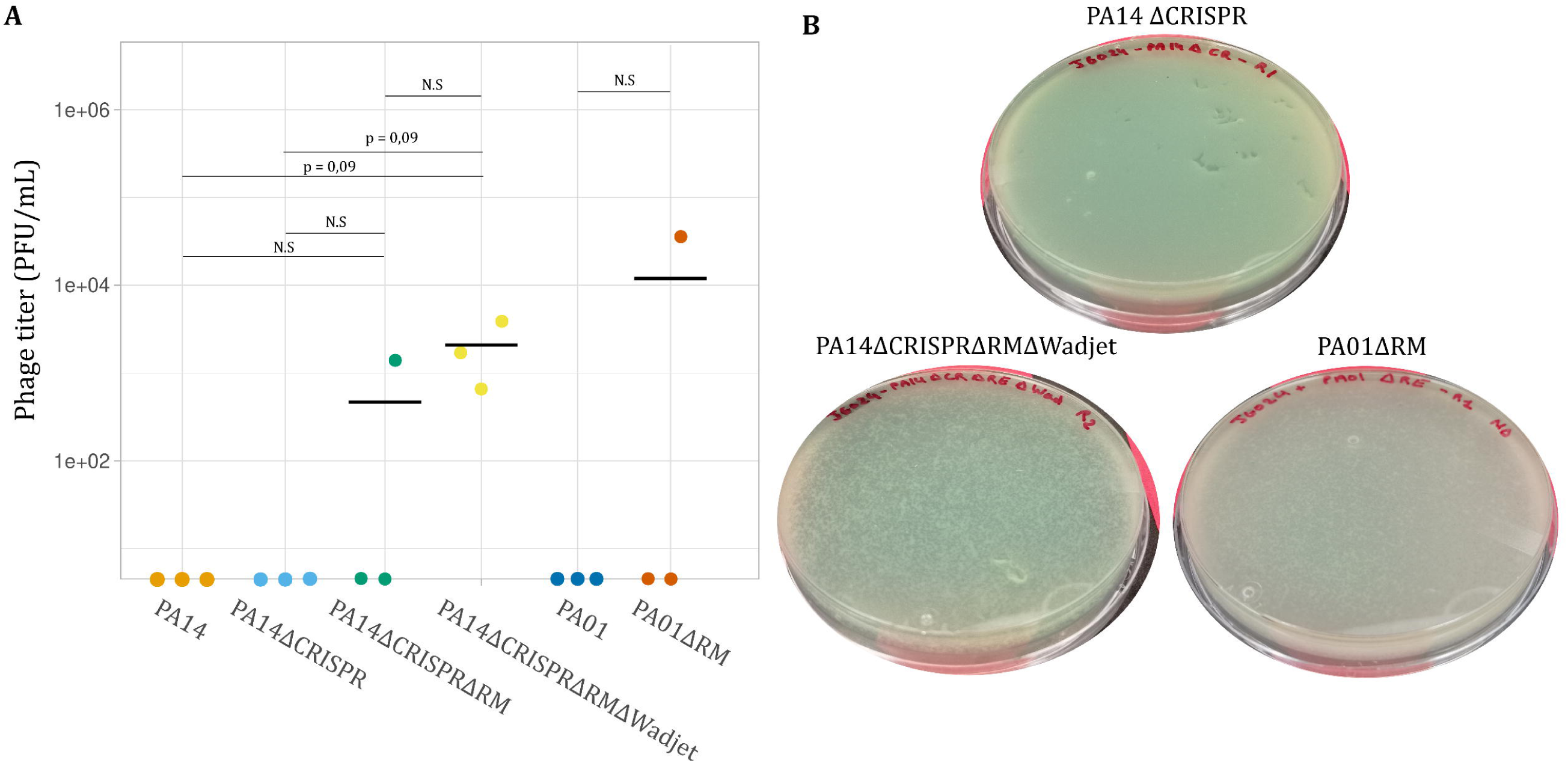
Phage reboot from yeast DNA in PA mutants. **A-** Reboot of linear JG024 phage DNA from yeast DNA in PA mutants. **B-** Example reboot result from yeast DNA obtained for linearized JG024 genomes in PA14 and PAO1 mutants.

### Phage genome validation by whole genome sequencing

Finally, we investigated whether the methodology created mutations in rebooted phages by performing whole genome sequencing on the wildtype DMS3 and JG024 phages and four clones of rebooted phages (two DMS3 reboots and two JG024 reboots) using Illumina NextSeq 6000 chemistry (Table S5, Figure S5). We expected two different types of mutations: stochastic mutations that appear during the phage replication process, which would be present unevenly across reads, and mutations linked to the methodology, which could be generated in yeast or during the cloning process. The latter should be represented as an ancestral mutation and thus be present on the overwhelming majority of reads, as a phage plaque is generated from a single phage that was generated in a single reboot event using a single copy of phage DNA produced in yeast. Comparing to the reference genome, we detected only 7 low-frequency mutations (not related to the methodology) in the two JG024 rebooted clones (details in SI-1), and no high-frequency mutations except the two already present in the WT. In the two rebooted DMS3 clones, we detected 13 low-frequency mutations (details in SI-1) and 8 high-frequency mutations, of which 7 were already present in the WT and only one was newly found in the two rebooted clones (position 36389, C to T). This last mutation is likely linked to the methodology since the site of this mutation is on a recombination arm used to add yeast elements during the cloning step. It is possible that the mutation was introduced due to heterogeneity in primers used to create the recombination arm. **Finally, we conclude that the methodology has high fidelity, and that mutations in the resulting phage can likely be controlled through stricter quality control of primers.**

### Expanding the methodology to other phages

To begin to explore how generalizable this method is, we tried to reboot two additional phages: vB_PaeP_PAO1_Ab05 ^58^ member of the Podoviridae family (Genus: Autographiviridae) and F8 phage^59^, a member of the Pbunavirus genus. In comparison with DMS3 and JG024, vB_PaeP_PAO1_Ab05 has a different genome structure with DTR (431 bp), and both vB_PaeP_PAO1_Ab05 and F8 are able to infect PAO1 but not PA14. We first successfully cloned vB_PaeP_PAO1_Ab05 and F8 genomes in yeast (Figure S6). Despite the inability of either phage to infect PA14, we successfully rebooted both in the triple-mutant strain PA14ΔCRISPRΔREΔWadjet. **This suggests that the methodology can be used to reboot diverse phages, even if the phage is not able to infect PA14.**

## Discussion

Using JG024, we developed a methodology for the construction of tailed PA phages, which are promising for phage therapy applications. This is the first step towards constructing “à la carte” phage genomes with specific traits and characteristics. Our analysis of JG024 has improved our understanding of this phage, particularly regarding the genome structure, with evidence indicating a circular permuted genome. The use of yeast as a platform for cloning and assembling phage genomes is an important step in advancing methodology for genomic manipulation of diverse phage, which must be coupled to a robust reboot strategy. We identified three major limitations to reboot from phage genomes cloned in yeast. By cloning and rebooting DMS3, another tailed phage, we demonstrated that different phage species have different reboot efficiencies. We identified bacterial defence systems that inhibit phage reboot from genome cloned in yeast. Finally, we demonstrate the possibility to reboot two more PA phages (vB_PaeP_PAO1_Ab05 and F8) that are not able to infect the strain use for reboot. Together, as a proof of concept, we demonstrate the possibility to reboot PA phages that belong to the three family of phages (Podoviridae, Siphoviridae, Myoviridae) and we identified barriers to the construction of synthetic, clinically-relevant phage.

In general, knowing the genome structure can influence the design of cloning and manipulation in yeast. For example, terminal ends could restrict the possible insertion sites for a yeast element. Our study suggests that the JG024 genome is circular permuted. Unicycler and Flye assembly suggest a circular genome in contrast to Trycycler assembly and Phageterm analysis. However, as Nextera transposon-based library preparation was used to prepare Illumina short-read sequencing data, it was expected that phage termini would not be detectable by methods such as PhageTerm because transposome sequence bias would likely misrepresent the distribution of read edge positions that are necessary for terminus prediction ^41, 60^. For example, Chung *et al.* (2017) ^60^ were unable to identify the termini of the novel *Bacillus cereus* phage SBP8a using Nextera-derived MiSeq sequencing data but identified a DTR of 2821 nt with Roche/454 sequencing data. Thus, the biased nucleotide frequency of the Nextera-derived reads may have altered the distribution of read edge positions to produce artificially high coverage regions, which were detected by PhageTerm as DTRs in this study. Indeed, Sanger sequencing results conflict with the DTR predicted by Trycycler and Phageterm. Furthermore, the successful amplification of overlapping fragments that cover the full JG024 genome, the digestion profile, and the mapping of long-read sequencing data (>30 000 bp) suggest that JG024 has a circular permuted genome. Experimental verification, e.g., by Southern blot analysis ^43^ is needed to make this observation conclusive. Further experiments could also verify the headful packaging strategy with a putative packaging site at position 59,376 bp.

Other characteristics, primarily related to transformation efficiency, are important for ensuring successful reboot (Figure 3, S2). As identified for at least 30% of tailed phage ^45^, we determined that JG024 is sensitive to chloroform. This is particularly important for experimental design as chloroform is used to release phage particles from bacterial cells for many types of phage experiments ^25, 61^. We also worked on transformation parameters that were already developed ^62, 63^ to obtain an optimized protocol for the reboot process for JG024 (Figure 3, S2). As this type of work expands, additional data will become available for more diverse phage. This will, in turn, enable generalized conclusions about the information needed to design and optimize a reboot protocol for any given phage.

Yeast has been extensively used as a cloning platform for high-length DNA molecules since 1980 ^64^. Different methods have been developed to clone genomic fragments and full-length virus or bacterial genomes in yeast ^47, 65–68^, and each of these methods requires the addition of yeast elements (Ars, Cen, Trp) to maintain the DNA molecule in yeast over time. These methods have allowed the cloning of genomes up to 1.8 Mb ^29, 66^, and as expected for small phage genomes, we successfully obtained several yeast clones containing stable JG024 genome and DMS3 genomes (Figure 3, 4). We used TAR-cloning ^65^ and genome assembly ^68^ methods to construct JG024 DNA, and those methods open up numerous possibilities for genome engineering during the cloning step. Furthermore, the yeast platform has the advantage of allowing the cloning of phage cargo genes that would be toxic for *E. coli* ^69^. In contrast to *E. coli* machinery that can recognize and express many prokaryotic genes ^69, 70^, the yeast machinery, which is eukaryotic, is unlikely to express prokaryotic genes, as most of the transcription signals are not recognized ^31^. This is particularly interesting for the cloning of phage genomes, which often contain genes for toxic proteins, such as Toxin-Antitoxin systems for phage selection pressure ^71^ or endolysin for phage release ^72, 73^. As observed in our study, JG024 genome cloning in *E. coli* was not functional. As *P. aeruginosa* and *E. coli* are closely related, we hypothesize that some JG024 genes can be expressed in *E. coli* and are toxic for the bacterial cell, but this still needs further experimental verification.

The yeast platform can also have several disadvantages. For example, as homologous recombination is efficient in yeast, even with the presence of yeast elements (ARS-CEN-Trp), DNA instability could occur through small DNA repeat sequences that can recombine and generate truncated versions of the genome over time ^74^. Our data showed that for JG204 phage, the genome is stable over 10 passages (Figure 4). The genome structure of phage containing DTRs could generate instability, but out results with vB_PaeP_PAO1_Ab05, in addition to a recent paper on *S. aureus* phages containing DTRs ^24^, suggest this is unlikely to be a widespread issue.

Another issue that we identified is the DNA methylation profile of the phage genome after production in yeast. DNA methylation in yeast is rare ^75, 76^, which is problematic for the use of this DNA to transform some bacterial strains. For example, it has already been described that for bacterial transplantation from genomes cloned in yeast, it was necessary to remove RM systems from the bacterial host strain or to perform in vitro methylation using cell extracts from the bacterial host strain ^28^. Indeed, the synthetic phage genome constructed in yeast is likely unmethylated and thus a target for cleavage by an RM system. PA possesses multiple antiviral defense systems, including CRISPR ^77^ and RM systems, and strains PAO1 and PA14 are no exceptions (Table S4). Our data confirm that RM can be problematic for DNA transformation from yeast DNA, particularly in PA14 where no phage reboot was observed in the presence of type-II RM genes (Figure 5D, 6A). We also see that the type-I RM system from PAO1, while not completely inhibiting DMS3 phage reboot, decreases the reboot efficacy (Figure 5D). This shows that different types of defense systems, in particular RM systems, will have different impacts on phage reboot, and removing those systems increases the probability of success. Several other defense systems have been identified in PAO1 and PA14, of which Wadjet systems are particularly notable. These defense systems, recently described in *Bacillus subtilis* and *P. aeruginosa*, recognize and cut DNA based on its topology, resulting in reduced transformation efficiency in *B. subtilis* ^57, 78^. Another study showed that Wadjet JetABCD systems restrict circular plasmids in *B. subtilis* but a linear plasmid evades restriction by *E. coli* JetABCD *in vivo* ^79^. When removing the Wadjet system from PA14, we did not observe an increase in plasmid transformation efficiency (Figure S4A) but we observe a potential implication of Wadjet system on the reboot consistency from linear JG024 DNA genome previously cloned in yeast, with replicable results obtained using the strains without Wadjet system (Figure 6A). Several experiments are needed to understand the exact implications of Wadjet systems on phage reboot from in yeast-cloned genomes and to describe the molecular mechanism of Wadjet restriction in *P. aeruginosa*.

The use of yeast as platform requires the use of a yeast element for circularization and maintenance of the phage genome in yeast. It is unclear if the presence of the yeast element on the phage genome can be problematic for subsequent reboot (SI-I). Previous phage reboot papers that use yeast or *E. coli* as a manipulation platform do not describe the release of the yeast element before phage reboot ^23–25, 32, 80^. However, *in vitro* genome assembly has been used to demonstrate that DNA circularity increased reboot efficiency ^80^. Using our reboot conditions, JG024 was not able to reboot as a circular molecule (SI-I). This is possibly due to DNA length, which is higher by 9.3 kb when containing yeast and *E. coli* elements, or topology rather than DNA circularity. More experiments are needed to understand this phenomenon and whether it impacts other phage reboot methodologies.

Finally, in this work, we have developed a method for rebooting clinically relevant *P. aeruginosa* phage. This is the first step towards important genome engineering that could be performed on JG024, DMS3 and other phages to improve and add specific phenotypic traits that could be useful for phage therapy applications. For example, changing the receptor to, e.g., expand the host range of PA strains that could be infected ^25^, adding anti-CRISPR proteins to prevent CRISPR adaptation by the targeted PA strain ^21^ or adding anti-quorum sensing proteins to inhibit biofilm production ^53^. This work thus represents a critical step towards using phage therapy to overcome antimicrobial resistance and treat infection.

## Materials and Methods

### Oligonucleotides and plasmids

All oligonucleotides used in this study were supplied by Integrated DNA (IDT) and are described in Table S1. All plasmids constructed and used in this study are listed in Table S2.

### Microbial strains and culture

*P. aeruginosa* strain PA14 (Tax ID: GCF_000404265.1) was provided by Dr. Egon Ozer (Northwestern University). *Pseudomonas* phage DSM 19871 (JG024) ^39^ was obtained from DSMZ (Braunschweig, Germany). Phage DMS3 ^50^, *Pseudomonas aeruginosa* PAO1 (Tax ID: NC_002516) and PA14ΔCRISPR ^51^ were provided by Pr George A. O’Toole (Geisel School of Medicine at Dartmouth). *P. aeruginosa* strains were cultivated at 37°C in Lysogeny Broth (LB) media or Vogel-Bonner minimal medium (VBMM). Gentamycin at 50 µg mL^-1^ or Carbenicillin at 300 µg mL^-1^ were used for selection. F8 phage ^59^ and vB_PaeP_PAO1_Ab05 phage ^58^ were provided by Pr Joseph Bondy-Denomy.

*S. cerevisiae* VL648-N was provided by Dr. Carole Lartigue (INRAE). *S. cerevisiae* MAV203 (Thermo Scientific, 11445012) and VL648-N were cultured in YPDA (Takara, 630464) or SD-Trp Broth (Takara, 630411 and 630413) at 30°C with shaking at 225 rpm.

### S. cerevisiae VL648-N transformation procedure

Phage genome cloning in non-commercial VL648-N strain was performed following ^49, 65^ and ^47^ with several modifications. For cloning half genome of JG024, *in vitro* cleavage of phage DNA was performed using the *Streptococcus pyogenes* CRISPR system. sgRNA was produced using EnGen® sgRNA Synthesis Kit (NEB, E3322S), primer D31 or D32, and purified using Monarch® RNA Cleanup Kit (NEB, T2040S). Cas9 nuclease (NEB, M0386S), sgRNA and 1 µg of phage DNA were incubated at 37°C for 20 min. Cas9 was then inactivated by incubation at 65°C for 10 min.

### Yeast DNA extraction

Individual yeast colonies were picked and streaked on SD-Trp and incubated 2 days at 30°C. Then, one isolated colony per streak was patched on SD-Trp plate and incubated for 2 days at 30°C. Total genomic DNA was extracted from yeast transformants according to ^65^.

### Phage reboot protocol

Phage reboot was performed using a previously described PA electroporation protocol with some modifications. Different parameters were tried as described in the results section. Finally, MgSO4 buffer was used for washing cells, 100 ng of phage DNA was used as control, LB was complemented with 1 mM MgSO4 and incubation of 3 to 24 h was performed for cell regeneration and phage production. For chloroform assays, 2-3 drops of chloroform were added to the cell suspension after incubation to kill bacterial cells and release the phages. For reboot from yeast DNA, separation and release of a linear phage DNA from the yeast recombination matrix was performed using 10 µg of in yeast DNA digested using SmaI (NEB, R0141S) for JG024 and ScaI (NEB, R3122S) for DMS3, vB_PaeP_PAO1_Ab05 and F8. Restriction enzymes were inactivated by 80°C heat inactivation for 20 min and DNA was then kept at 4°C until transformation.

To quantify PFUs after phage reboot incubation, serial dilutions of supernatant were made with LB media. 300 µL of supernatant were separately mixed with 200 µL of mid-exponential PA14 cells and 4 mL of LB soft agar (0.8%) complemented with 1 mM MgSO4 and prewarmed to 55°C. The agar mixture was then poured onto LB plates, incubated overnight at 37°C. Plates containing phage plaques were then counted.

### Genome sequencing analysis

All genome sequencing was sent to the Microbial Genome Sequencing Center (MiGS; Pittsburgh, PA) for library preparation, short-and long-read sequencing (Illumina and Oxford Nanopore technologies [ONT], respectively), and de novo assembly. Short reads were obtained using the Illumina DNA Prep Kit, IDT 10bp UDI indices, and the Illumina NextSeq 2000 platform ^81^. Illumina paired-end reads (2 x 151 bp) was provided as fastq files. For *P. aeruginosa* strains verifications, analyses were made using Galaxy (https://usegalaxy.eu/) and are presented in Table S5. Illumina reads were trimmed using Trimmomatic (V 0.38.1; Sliding Window 10, 20 ; Drop read below Minimal length 150), mapped using BWA-MEM (V 0.7.17.1), Samtools sort (V 2.0.3), MPileup (V 2.1.1), and variants were detected using VarScan mpileup (V 2.4.3.1; Minimum coverage 20, Minimum supporting read 15, Minimum Base quality 20, Minimum variant allele frequency 0.5, Minimum homozygous variants 0.75). For defense systems mutants, deletions were verified using JBrowse (V 1.16.11). For DMS3, JG024 phage and rebooted clone verification, Illumina NextSeq 6000 sequencing was performed, presented in Table S5. Demultiplexing, quality control and adapter trimming was performed by MiGS with bcl-convert (v4.0.3). Quality control was checked using FastQC (v0.11.5) (https://www.bioinformatics.babraham.ac.uk/projects/fastqc/) and MultiQC (v1.11). To detect mutations, read mapping was performed using BWA-MEM (v0.7.12) with default parameters. DMS3 and JG024 WT and clones were mapped to their NCBI reference genomes, NC_008717 and NC_017674, respectively. Mapped reads were converted to BAM format using the Samtools (v1.6) view command, sorted using the sort command and reads were piled using the mpileup command. Indels and SNPs were identified using VarScan (v2.4.6) set to a --min-coverage=30, --min-reads2=20, --min-var-freq=0.01 and --min-freq-for-hom=0.75. All other VarScan parameters were run as default. VCF outputs from VarScan were visualized using IGV (v.2.8.10). Specific SNP and indel locations were compared with reference genome annotations on NCBI.

In addition, for JG024 WT, ONT sequencing were prepared using Oxford Nanopore’s “Genomic DNA by Ligation” kit (Oxford Nanopore Technologies, Oxford, UK) and sequenced on a MinION R9 flow cell. ONT long reads were provided as fastq files. Assembly using Flye (V 2.6) was performed using Nanopore reads filtered with Filter FASTQ (V 1.1.5, Minimum size 30 000 bp). The fasta assembly file was polished using 4 rounds of Pilon (1.20.1) combined with Illumina short reads. For Figure 2C observation, Nanopore reads were filtered using Filter FASTQ (V 1.1.5, Minimum size 30 000 bp), mapped using BWA-MEM (V 0.7.17.1) and observed on IGV ^82^.

## Supporting information

See Supplementary file

## Acknowledgements

Acknowledgements and funding sources

DMS3 phage, pMQ30 plasmid and PA14ΔCRISPR, were kindly provided by Pr George O’Toole. D3 phage, F8 phage and vB_PaeP_PAO1_Ab05 phage were kindly provided by Pr Joseph Bondy-Denomy. Yeast strain VL6-48N and PCC1-YTrp plasmid were kindly provided by Dr Carole Lartigue. Thanks to Mustafa Ismail for help during experiments. This work was funded in part by the Northwestern University McCormick School of Engineering Research Catalyst Program, the Walder Foundation (Innovation Top-Up Award; Hartmann AGMT 12/16/21), The National Science Foundation Graduate Research Fellowship (Grant Number DGE-2234667), and the National Institutes of Health’s National Center for Advancing Translational Sciences (Grant Number TL1TR001423). The content is solely the responsibility of the authors and does not necessarily represent the official views of the National Institutes of Health or any other funding agency.

## Author contribution

*T.I and R.R contributed equally to the work. T.I, R.R, and E-M.H designed the experiments; T.I and R.R performed the experiments; T.I, C.W, S.H, and E-M.H. analyzed the data and results; T.I, E.O and E-M.H wrote and improved the manuscript.

## Notes

The authors declare no competing financial interest.

---

**Figure S1:**
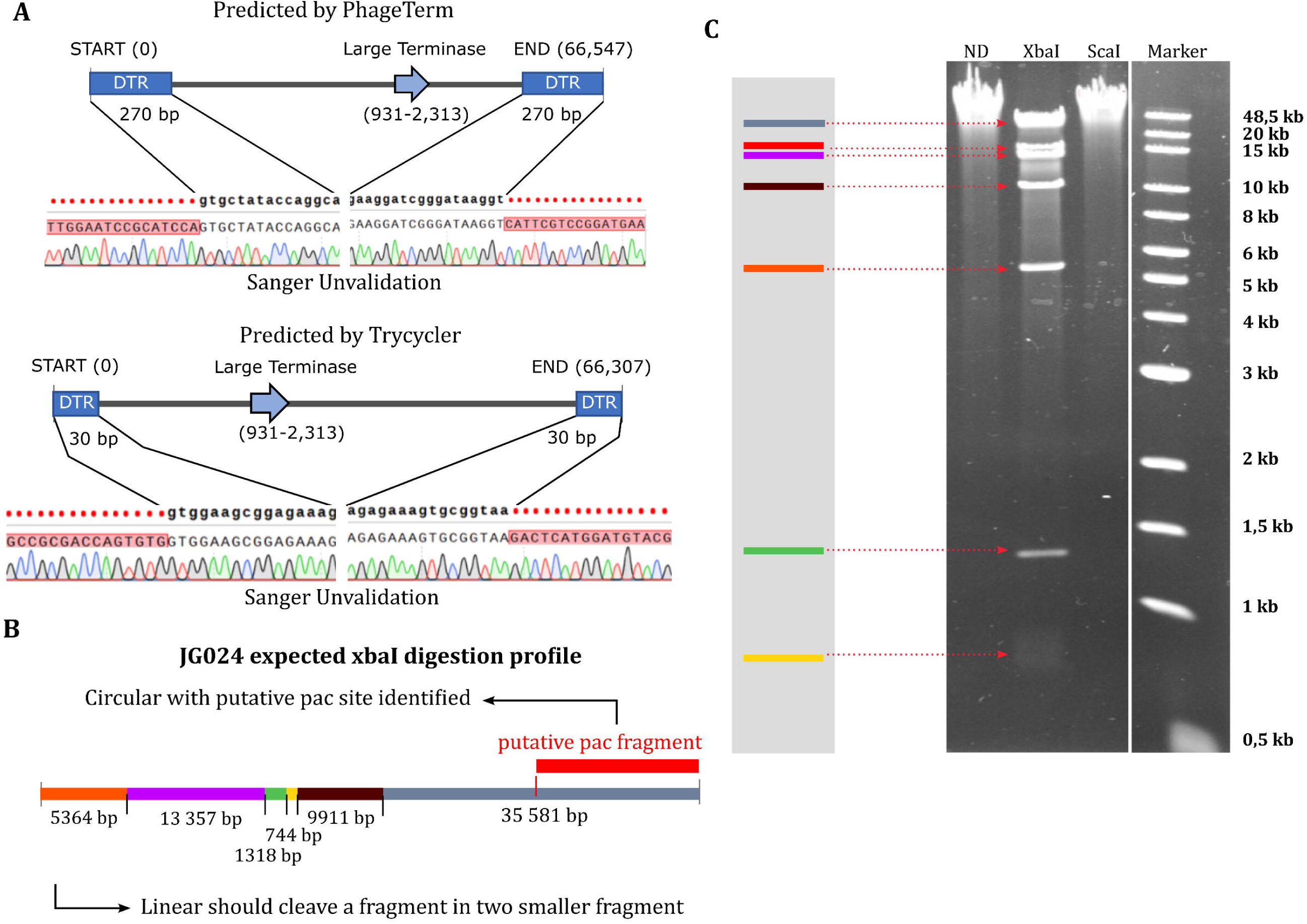
Invalidation of JG024 linear genome. **A-** Sanger sequencing of the two-region predicted to be JG024 DTR**. B,C-** Simulation of expected XbaI digestion profile according to circular or linear form and comparison to the obtained profile.

**Figure S2:**
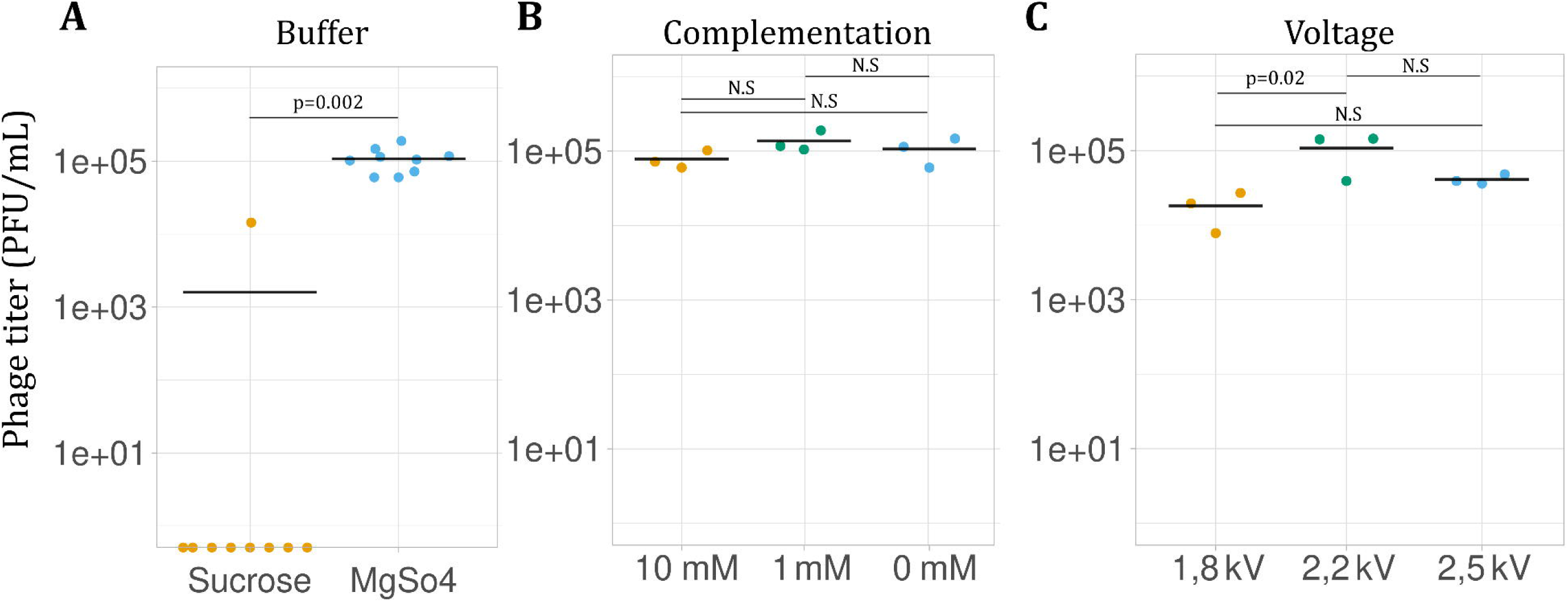
Other parameters tested for reboot protocol. Graphical representation for all parameters tested: **A-** Buffer used for washing PA cells (Sucrose 300 mM or MgSO4 1 mM), **B**Media complementation with MgSO4 (0, 1, 10 mM) and **C-** Voltage used for electroporation.

**Figure S3:**
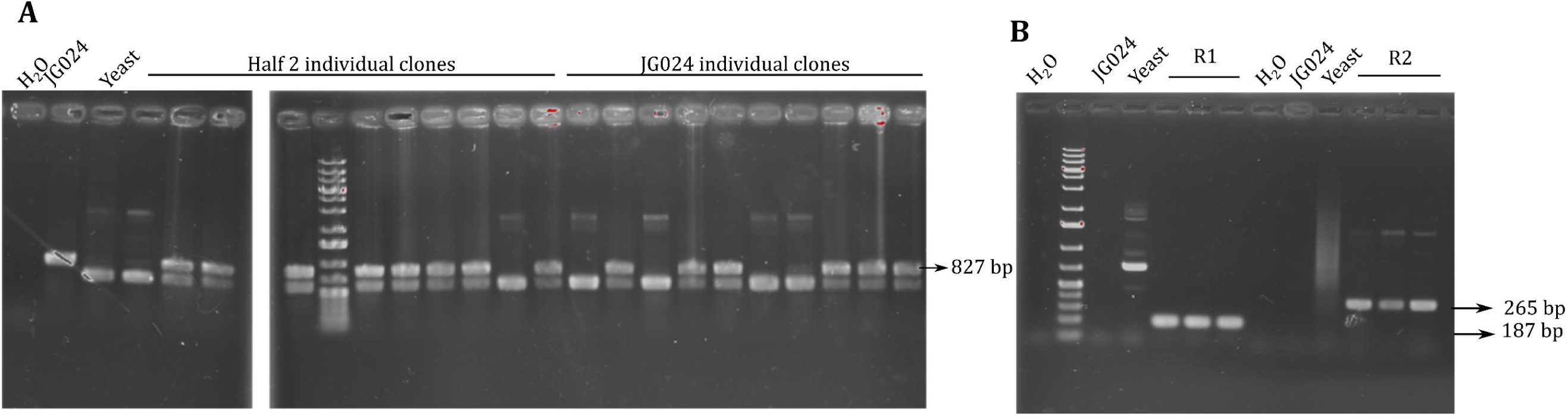
Screening of JG024 in yeast clones. **A-** Example of simplex PCR performed to validate first yeast clone screening. **B-** Example of recombination PCR to amplify recombination scars.

**Figure S4:**
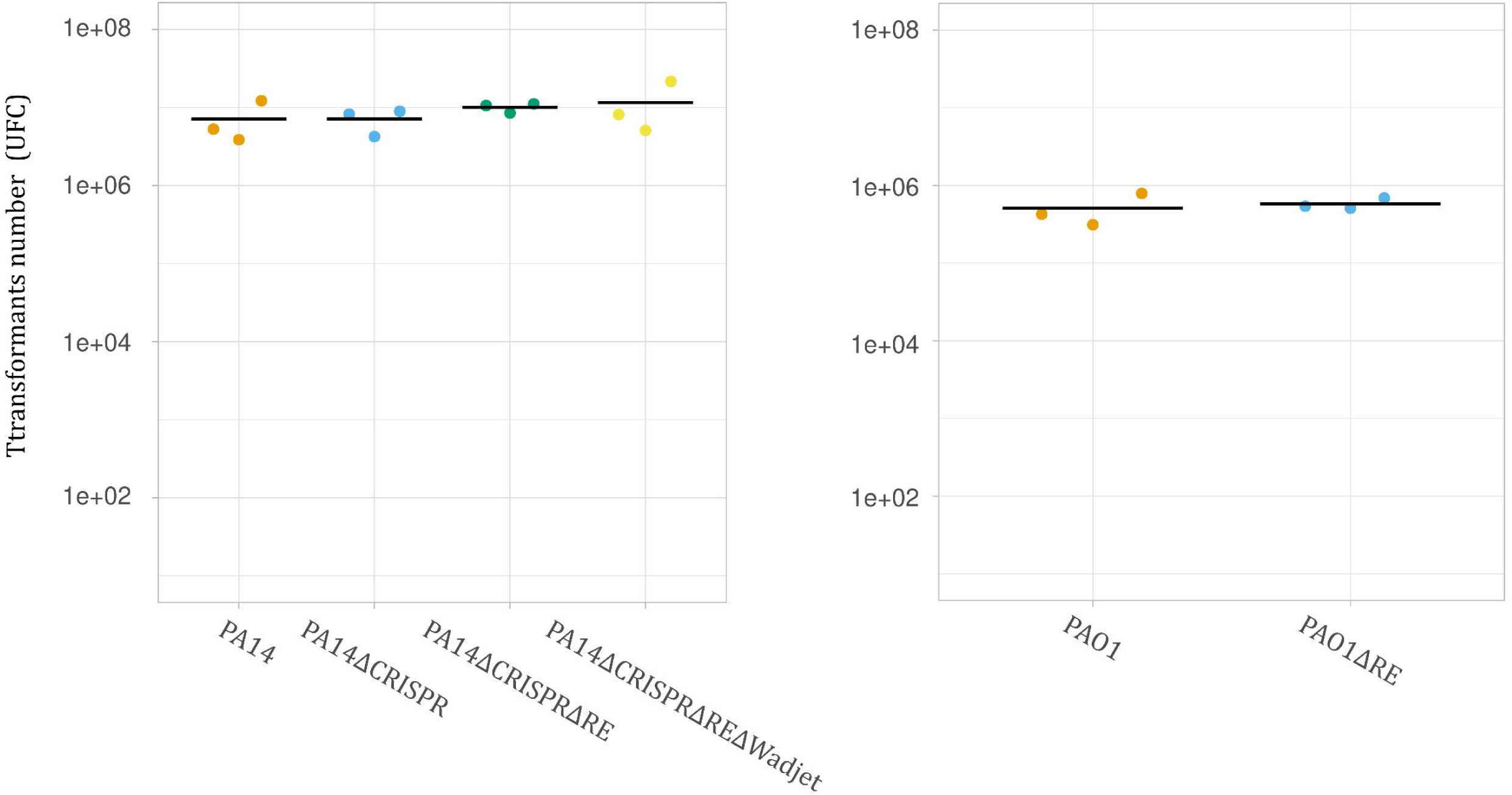
Validation of transformation and reboot in PA mutants. **A-** Transformants obtained in each PA mutant strain after pucP-20T-tsapphire transformation. **B-** Reboot efficiency obtained in each PA mutant strain after reboot using DMS3 genome.

**Figure S5:**
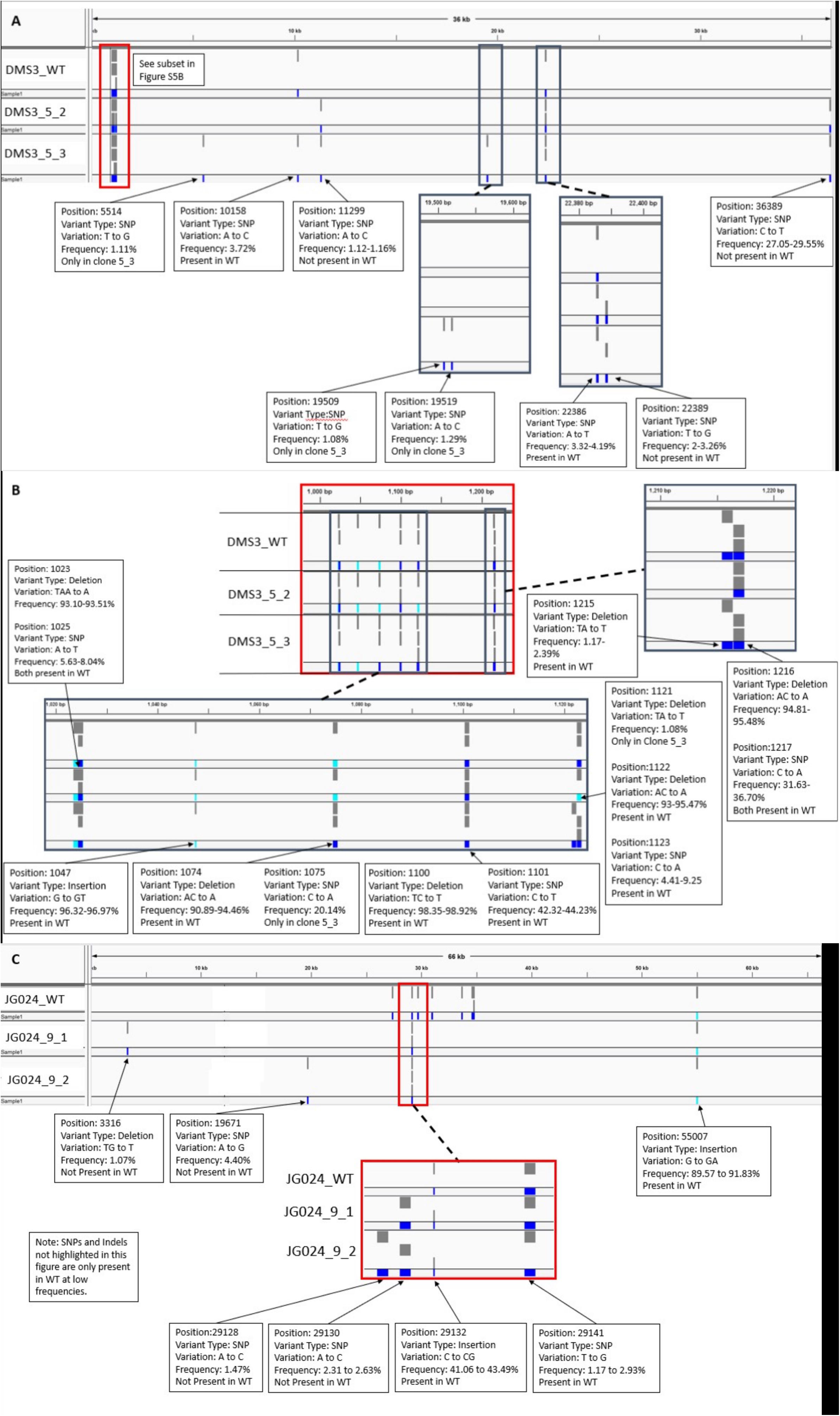
Sequencing analysis on phage rebooted clones. **A-** Visualization of DMS3 WT and clone variations compared against the NCBI reference genome. **B-** Subset of DMS3 genome positions 1000 – 1250 to show region variants. **C**-Visualization of JG024 WT and clone variations compared against the NCBI reference genome. Mutations not referenced on either visualization were only found in the WT and can be referenced in Table S5.

**Figure S6:**
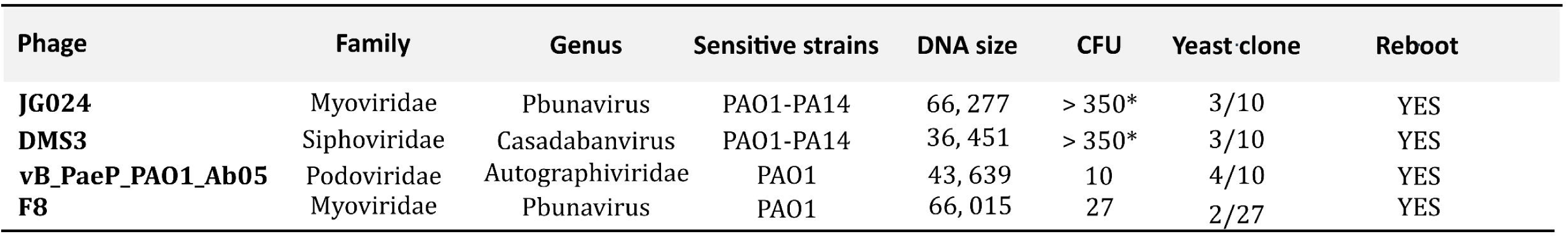
Summary table of phage reboot.

