## Supplementary file for "A synthetic biology approach to assemble and reboot clinically-relevant *Pseudomonas aeruginosa* tailed phages"

### Supplementary Information

#### Supplementary text

##### SI-1. Results

###### Rebooting circular versus linear genome.

The phage genome extracted from yeast clones is circular and contains yeast elements, along with *E. coli* elements for JG024 and DMS3. Two restriction sites allow the removal of these elements, resulting in the release of a linear phage genome. In our experiments, we attempted to reboot both forms of phage DNA: the circular form containing yeast elements and the linear form free from yeast elements. Interestingly, we observed that by using linear phage DNA, we were able to successfully reboot all tested phages, including JG024, DMS3, F8, and vB_PaeP_PAO1_Ab05. However, under our specific conditions, JG024 was unable to be rebooted when in its circular form. This observation highlights a lack of understanding regarding the reboot process, which may limit the effectiveness of the technique for certain phages.

###### Sequencing analysis

We sequenced the DMS3 WT phage that was initially used for reboots alongside two DMS3 reboots, DMS3 5_2 and DMS3 5_3. Sequencing coverage was 49 889x, 35 741x, and 33 377x for DMS3 WT, DMS3 5_2 and DMS3 5_3, respectively. DMS3 WT was found to have 13 variant positions: 7 SNPs and 6 indel mutations. DMS3 5_2 also had 13 variant positions: 6 SNPs and 7 indel mutations. DMS 5_3 had 21 variant positions: 13 SNPs and 8 indels. A total of 21 unique SNP or indel mutations were found across the three phages. Ten of the mutations occur in all three sequenced DMS3 phages and at similar frequencies, suggesting that these variations existed within our WT population prior to reboot. Three mutations occur in the WT and one clone but not the other, again at similar frequencies. Eight total mutations occur that are not represented in the WT sequencing effort, six of which occur at a frequency of 3.26% or less. One SNP, a C to an A, occurs at position 1075 in DMS3 5_3 at a frequency of 20.14%. In context of other variations, a deletion of a C occurs at position 1074 in all three genomes at frequency of 90.89% or higher. In the NCBI RefSeq DMS3 genome, starting at position 1074, the genome reads A-A-C-C-C. In our DMS3 WT the deletion of a C at this position is the standard, giving us A-A-C-C. While the 1075 SNP A to C in only DMS_5_3 reads A-C-C-C, returning that run of three C nucleotides, suggesting that this behavior is implicit to the genome rather than caused by methodology. Especially as both variations occur in a non-coding region of the phage. At position 36389, a C to T substitution occurs at a frequency of 27.05-29.55% in the rebooted phages but not our

WT DMS3. This variation occurs in the recombination arms and is the only one that is probably linked to the methodology.

The two JG024 reboots, JG024 9_1 and JG024 9_2, had sequencing coverage of 18 760x and 19 773x respectively using short read chemistry. The JG024 WT was sequenced at an earlier point and had 13 variant positions: 11 SNPs with low frequency detections and 2 insertions present in most of the population. JG024 9_1 had 5 variant positions: 2 SNPs and 3 indels and JG024 9_2 had 6 variant positions: 4 SNPs and 2 insertions. Eleven variations occur in the WT but neither clone, at a range of 1.01 to 23.19% frequency. Since neither clone has any of these mutations, they were most likely not present in the yeast parental lineage. A total of 7 unique variant positions occurs between the two clones. Four of these mutations occur in only one of or both of the clones, but are not present in the JG024 WT. All four of these mutations occur at a frequency of 4.40% or less. The other three mutations identified in both clones were shared at similar frequencies with the JG024 WT. **Sequencing and variant analysis demonstrate that the method used for reboot is unlikely to create mutations, the only one identified in DMS3 rebooted clones were not deleterious mutations that significantly impact the structure or function of the rebooted phage particles.**

#### SI-2. Materials and Methods

**Phage DNA extraction.** Liquid lysates of phage were made according to [1] by infecting 50 mL of mid-exponential PA14 cells in LB broth with 50 µL of phage lysate and incubating the cultures at 37°C overnight with shaking at 180 rpm until clear. Prior to DNA extraction, 24 mL of phage lysate was concentrated using polyethylene glycol (PEG) precipitation as described by [2] with modification. Briefly, aliquots of 1.2 mL of filter-sterilized phage lysate were treated with 300 µL of phage precipitation solution (20% (*w/v*) PEG 8,000 [MilliporeSigma, Burlington, MA, USA], 2.5 M NaCl). After gentle mixing and overnight incubation at 4°C, phage particles were precipitated by centrifugation at 13,000 x *g* for 30 min at 4°C. The supernatant was removed, and the visible phage pellet was resuspended in 120 µL of SM buffer (100 mM NaCl, 8 mM $MgSO_4 \cdot 7H_20$, 50 mM Tris-HCl [pH 7.4]). The PEG-concentrated phage lysate in SM buffer was then treated with DNase I to remove bacterial host genomic DNA by incubating samples of 360 µL of PEG-concentrated phage lysate, 40 µL of 10x DNase I (New England Biolabs, Beverly, MA, USA), and 1 µL of DNase I (2000 U/mL, New England Biolabs, Beverly, MA, USA) at 37°C for 30 min. To inactivate the DNase I, 10 µL of 0.5 M EDTA was added to each sample prior to heat inactivation at 75°C for 10 min. Phage DNA was then extracted from the processed lysate using a NucleoBond HMW DNA kit (Machery-Nagel, Duren, Germany) according to manufacturer's instructions with minor modification. The Proteinase K treatment was extended to an overnight incubation at 50°C. Furthermore, 20 µL of 5 mg/mL linear acrylamide (GeneLink, Orlando, FL, USA) was added to 5 mL of the eluted DNA in Buffer H5 prior to DNA precipitation. DNA precipitation was conducted by adding an equal volume of room-temperature isopropanol to the sample prior to centrifugation at 15,000 x *g* for 30 min at 4°C. After discarding the supernatant, the DNA pellet was resuspended in 200 µL of resuspension buffer (5 mM Tris-Cl [pH 8.5], Machery-Nagel, Duren, Germany). Prior to sequencing and downstream analysis, the resuspended DNA was desalted using agarose cones as described by [3].

**Plasmid construction for PA deletions.** To prepare plasmid for deletions in PA, the backbone shuttle vector pMQ30 [4] was used. Primers A1/A2, B11/B12 and C21/C22 were used and the Q5® High-Fidelity DNA Polymerase (M0491) kit to amplify the desired plasmid region. Primers A3/A4 and A5/A6, B13/B14 and B15/B16, C23/C24 and C25/C26 to amplify 1 kb flanking DNA of the target site in the PA genome, respectively for deletion of Type II restriction system (CIA_03834 and CIA_03831) and Wadjet system (CIA_04813, CIA_04814, CIA_04815, and CIA_04816) in PA14 strain and Type I restriction system (NP_251425.1 and NP_251424.1) in PAO1 strain. PCR products were submitted to restriction with the DpnI enzyme (NEB, R0176S) following the manufacturer's protocol. Both products were purified using GFX™ PCR DNA or Gel Band Purification Kit (GE-healthcare). The NEBuilder HiFi DNA Assembly Cloning Kit (NEB, E5520S) was used to assemble the fragments into pMQ30ΔRE-PA14, pMQ30ΔWadjet and pMQ30ΔRE-PAO1. Two µL of the assembly was transformed into *E. coli* NEB 5-alpha (NEB, C2987H). Transformants were screened after DNA minipreps using NucleoSpin Plasmid kit (Macherey-nagel, 740588.50) and enzymatic digestion.

***Pseudomonas aeruginosa transformation procedure*.** Electrocompetent PA cells were prepared according to [5,6] with minor modification. Briefly, one colony of PA was inoculated into 6 mL LB medium, grown overnight at 42°C with 100 rpm shaking until stationary phase (~$1.8 \times 10^8$ CFU $mL^{-1}$), and pelleted at 4,500 x *g* for 10 min at room temperature. The cell pellet was then washed twice with 1 mL of 1 mM room temperature $MgSO_4$ before final resuspension in 100 µL of 1 mM room temperature $MgSO_4$. 100 µL of electrocompetent cells were mixed with 100ng of plasmid and the mixture was transferred to a 2 mm gap electroporation cuvette. A 2.2 kV pulse was applied with a MicroPulser (Bio-Rad, Hercules, CA, USA) and 950 µL of LB medium was added immediately to the cuvette. The cell suspension was then quickly transferred to a 17 mm x 100 mm culture tube and incubated at standard culture conditions for 2 hrs. Cells were diluted and plate in LB with antibiotic.

PA mutants were produced using conjugation protocol according to [7,8]. After several passages (2-3), mutants were screened using A7/A8, C27/C28 and B17/B18 respectively to amplify scars on restriction-modification deletion and Wadjet deletion in PA14 and restriction-modification deletion in PAO1. A9/A10, B19/B20 and C29/C30 were used to confirm counter selection and loss of WT genotype.

**Yeast recombination matrix preparation.** For the synthetic phage construct, the yeast artificial chromosome component (ARS, CEN, Trp) from pYES1L-URA was linearized by digestion with BamHI and EcoRI and gel-purified using the Zymoclean Large Fragment DNA Recovery kit (Zymo Research, Irvine, CA, USA). PCR products which spanned the JG024 genome were prepared with Platinum SuperFi II DNA polymerase (Thermo Scientific, Waltham, MA, USA) according to primer sets listed in Table S1. To facilitate yeast gap repair cloning, 50 bp homologous bases between the linearized YAC and the JG024 genome were added to the first and last JG024 genomic fragments. Standard PCR conditions resulted in nine PCR products (G79 to G96) ranging from 2.2 – 8.0 kbp while long PCR resulted in three PCR products ranging from 16.0 – 32.0 kbp. All the PCR fragments were gel-purified using either the NucleoSpin Gel and PCR Clean-up Kit (Machery-Nagel, Duren, Germany) or the Zymoclean Large Fragment DNA Recovery kit (Zymo Research, Irvine, CA, USA).

For TAR-cloning from HMW extracted phage DNA (JG024 and DMS3), the yeast artificial chromosome component (ARS, CEN, Trp) were amplified with primers E53/E54, E53/E61, E59/E54, F75/F76, I105/I106 and J109/J110 that contain 60bp of homology with JG024, Half1 JG024, Half2 JG024, DMS3, F8 and vB_PaeP_PAO1_Ab05 phage ends respectively, using Advantage 2 polymerase (Takara, 639282). During primers design, blunt restriction site not find in JG024 or DMS3 genome were added for yeast artificial chromosome component elimination before phage reboot. To eliminate plasmid and avoid yeast transformant background, PCR products were submitted to restriction with the DpnI enzyme (NEB, R0176S) following the manufacturer's protocol. Both products were purified using GFX™ PCR DNA or Gel Band Purification Kit (GE-healthcare).

***S. cerevisiae MAV203 transformation procedure*.** To assemble the synthetic JG024 genome, the three long PCR products and the linearized YAC (200 ng of each PCR product and 100 ng linearized YAC in 10 µL water) were transformed into competent MAV203 yeast cells using the GeneArt High-Order Genetic Assembly System (Thermo Scientific, Waltham, MA, USA) according to manufacturer's instructions.

Yeast transformants were selected on a minimal synthetic defined medium without tryptophan (SD-Trp) (Takara, San Jose, CA, USA) agar plates at 30 °C for 3 days.

**Phage DNA screening in yeast.** Yeast clones were screened for both the presence of the phage genome and the correct assembly with recombination matrix by PCR, using the Advantage 2 Polymerase kit (Clontech) and specific primers located on either side of the target locus. Yeast transformants were then screened for phage genome completeness by multiplex PCR using specific sets of PCR primers for each phage (**Table S1**). Each set used several pairs of primers distributed across the bacterial genomes allowing the simultaneous amplification of fragments ranging from ~100 to ~1000 bp. The multiplex PCR was performed using the Qiagen Multiplex PCR Kit (Qiagen, 206143) according to the manufacturer's instructions.

#### SI-3. Supplemental references
